# Volatile-suppressed peptide signaling enhances volatile responses in plant-plant interactions

**DOI:** 10.1101/2025.01.20.634011

**Authors:** Lei Wang, Sara Hoefer, Pedro Jimenez-Sandoval, Hao Yu, Roxane Spiegelhalder, Jamie Waterman, Luke Hurni, Lingfei Hu, Lei Liu, David Jackson, Michael Raissig, Matthias Erb

**Affiliations:** Institute of Plant Sciences, University of Bern, Altenbergrain 21, 3013 Bern, Switzerland; Yazhouwan National laboratory, Sanya, Hainan 572024, P.R. China; The Plant Signaling Mechanisms Laboratory, Department of Plant Molecular Biology, University of Lausanne, 1015 Lausanne, Switzerland; Discipline of Botany, School of Natural Sciences, Trinity College Dublin, Dublin 2, Ireland; Institute of Soil and Water Resources and Environmental Science, College of Environmental and Resource Sciences, Zhejiang University, Hangzhou 310058, China; Cold Spring Harbor Laboratory, Cold Spring Harbor, NY 11724, USA; Oeschger Centre for Climate Change Research, University of Bern, Hochschulstr. 4, 3012 Bern, Switzerland

**Keywords:** maize, peptide signaling, plant volatiles, stomata, volatile emissions

## Abstract

Plant volatiles shape plant-plant interactions by acting as defense regulators and response factors. While plant volatile biosynthesis is well understood, how their emission is regulated remains largely elusive. Here, we show that small peptide signaling regulates induced volatile release in maize. Following herbivore attack, green leaf volatiles such as (*Z*)-3-hexenyl acetate (HAC) are released and induce terpene and indole emissions from neighboring plants. This process is accompanied by reduced expression of the ZmCLE1E9 gene and the ZmBAM1A, ZmBAM1B and ZmBAM3C receptor genes in HAC-exposed plants. Exogenous ZmCLE1E9 peptide inhibits HAC-triggered volatile release by limiting stomatal aperture. This inhibition disappears in the *Zmbam1a/Zmbam1b/Zmbam3c* triple mutant. Molecular docking supports ZmCLE1E9 and ZmBAMs as ligand-receptor pairs. Furthermore, *Zmcle1e9* and *Zmbams* triple mutants show increased volatile emissions upon HAC exposure. In summary, we show that upon HAC perception, maize plants enhance their capacity to release terpenes and indole via the suppression of CLE1E9 signaling. This behavior allows maize plants to rapidly deploy volatile cues in response to stress volatiles and thus shape the infochemical dynamics of multitrophic environments.

## Introduction

Plants use large quantities of carbon fixed by photosynthesis to produce volatiles. These volatiles, after emissions into the atmosphere, shape plant interactions with the environment (Karban, 2021; Zhou 周绍群 & Jander, 2022). Herbivore-induced plant volatiles (HIPVs) function in complex ways regulating the interactions between plant-plant, plant-herbivore and plant-natural enemy (Turlings & Erb, 2018). Many plants rapidly emit green leaf volatiles (GLVs) upon fresh herbivore damage. These volatiles can induce defense in neighboring plants, deter herbivorous insects and attract natural enemies (Matsui & Engelberth, 2022). HIPVs that are released later include terpenoids and aromatic compounds such as indole. These volatiles can attract natural enemies and thereby protect plants, and can influence herbivore behavior directly by acting both as repellents and attractants (Turlings & Erb, 2018).

GLVs are derived from the membrane lipid of damaged plant cells. Their production and emission happen within minutes upon attack from herbivorous insects or simply mechanical damage (Tanaka *et al*, 2018; D’Auria *et al*, 2007; Matsui & Engelberth, 2022). Other HIPVs typically require de novo biosynthesis upon herbivory or induction by molecules associated with herbivory (Turlings & Erb, 2018). In maize, molecular patterns identified from insect regurgitant, such as volicitin and caeliferin, have been reported to trigger increased emission of indole, monoterpenes, sesquiterpenes and the homoterpenes (*E*)-4,8-dimethyl-1,3,7-nonatriene (DMNT) and (*E,E*)-4,8,12-Trimethyl-1,3,7,11-tridecatetraene (TMTT) (Alborn *et al*, 1997; Alborn *et al*, 2007; Hu *et al*, 2019; Wang *et al*, 2023). The elevated volatile emissions are also reflected at the transcriptional level, shown by the increased expression of volatile biosynthesis genes such as *ZmIGL*, *ZmFPPS3*, *ZmTPS2*, *ZmTPS10* and *ZmCYP19* (Hu et al, 2019). Interestingly, GLVs from *Spodoptera exigua*-attacked maize plants can trigger increased volatile production and emissions in the neighboring plants which are not experiencing herbivory (Engelberth *et al*, 2004). While the regulation of volatile biosynthesis is well understood, little is known about the molecular regulation of transport and emission.

Stomata are the major release sites of foliar volatiles (Lin *et al*, 2021b). Therefore, stomatal activity greatly affects volatile emissions. In maize, both dark exposure at midday and abscisic acid(ABA) treatment can cause stomatal closure to reduce elicitor-triggered farnesene emissions (Seidl-Adams *et al*, 2015). Similarly, stomatal closure caused by the *Helicoverpa zea* glucose oxidase can reduce volatile emissions in tomato (Lin *et al*, 2021a). Stomatal control is tightly regulated. For instance, herbivory by chewing insects can cause stomatal closure, most likely as a means to reduce local water loss (Nabity *et al*, 2013; Meza-Canales *et al*, 2017). How plants balance the need to close stomata to avoid water loss from the damaged leaves and the need to keep stomata open to emit HIPVs to execute their ecological functions remains an intriguing question.

Plant peptide hormones are important signals regulating various aspects of plant development and responses towards environmental stimuli (Matsubayashi, 2014; Rzemieniewski & Stegmann, 2022). Several families of plant peptides and their corresponding receptors have been reported to regulate stomatal immunity. In *Arabidopsis*, the plant elicitor peptide (Pep) family members AtPep1 and AtPep2 close stomata to limit the entry of bacterial pathogens, next to triggering other anti-pathogen defenses (Huffaker *et al*, 2013; Zheng *et al*, 2018). The maize peptide ZmPep3 triggers anti-herbivore defense and increased emission of volatiles (Huffaker *et al*, 2013), though the role of stomata in this process is unclear. Recently, a group of newly identified plant peptides called SCREWs were reported to reopen stomata that are closed in anti-pathogen defense (Liu *et al*, 2022). The CLAVATA3/ESR-RELATED (CLE) family is another peptide family that regulates stomatal activity or stomatal development. AtCLE9 and AtCLE10 can customize stomatal production by binding to the receptors AtHSL1 or AtBAM1 (BARELY ANY MERISTEM1) (Qian *et al*, 2018; Vatén *et al*, 2018; Roman *et al*, 2022). Interestingly, AtCLE9 also functions in guard cells to close stomata (Zhang *et al*, 2019). Another peptide AtCLE25, induced by dehydration in the roots, travels to the leaves to trigger stomatal closure through the BAM receptors in *Arabidopsis* (Takahashi *et al*, 2018). Maize genome contains dozens of genes encoding CLE peptide precursors (Je *et al*, 2016; Goad *et al*, 2017). Whether CLE peptide signaling is involved in regulating induced volatile emissions in maize is unclear.

Here, we investigated how maize plants regulate their volatile release in response to the wound- and herbivory induced volatile (Z)-3-hexenyl acetate (HAC). We find that upon HAC perception, maize plants suppress ZmCLE1E9 signaling through ZmBAM1A, ZmBAM1B and ZmBAM3C to boost their volatile release, most likely by keeping stomata open. We thus uncover a molecular and physiological mechanism that allows plants to rapidly emit volatiles in response to stress volatiles and thereby contribute to chemical interactions between plant populations and their environment.

## Results

### HAC exposure suppresses CLE signaling

We recently reported that immature leaves are the major volatile sensing organs in maize. They respond strongly to herbivory induced volatiles and HAC exposure on the transcriptional and hormonal level, resulting in the release of terpenes and indole (Wang *et al*, 2023). To identify novel players that may regulate volatile-induced foliar volatile emissions, including peptide hormones, we re-mined the RNA-seq data with a focus on CLE peptides and their potential receptors. In both immature leaves of V4 maize plants, the expression of the ZmCLE1E9 precursor gene and expression of *ZmBAM1B* were downregulated by HAC. In leaf 6, the youngest leaf of V4 maize plants, the expression of *ZmBAM1A* and *ZmBAM3C* were also downregulated by HAC (Figure 1A and 1B). To gain deeper insights into the expression pattern of the *CLE* and *BAM* genes, we traced their transcript level in the immature leaf 3 from intact V2 stage B73 maize plants. The expression of *ZmCLE1E9*, *ZmBAM1A, ZmBAM1B* and *ZmBAM3C* was downregulated half an hour after HAC exposure. *ZmCLE1E9*, *ZmBAM1A* and *ZmBAM1B* transcripts reached the lowest level after one hour, before returning to the basal level (Figure 1C, 1D). We also observed reduced expression of the ZmCLE1B3 precursor gene at the half hour point (Figure 1D). The expression pattern of *ZmCLE1E9* was particularly interesting. In a HAC exposure time series experiment, its downregulation mirrored the upregulation of volatile biosynthesis genes (Figure 1D, Figure EV1).

**Figure 1.**
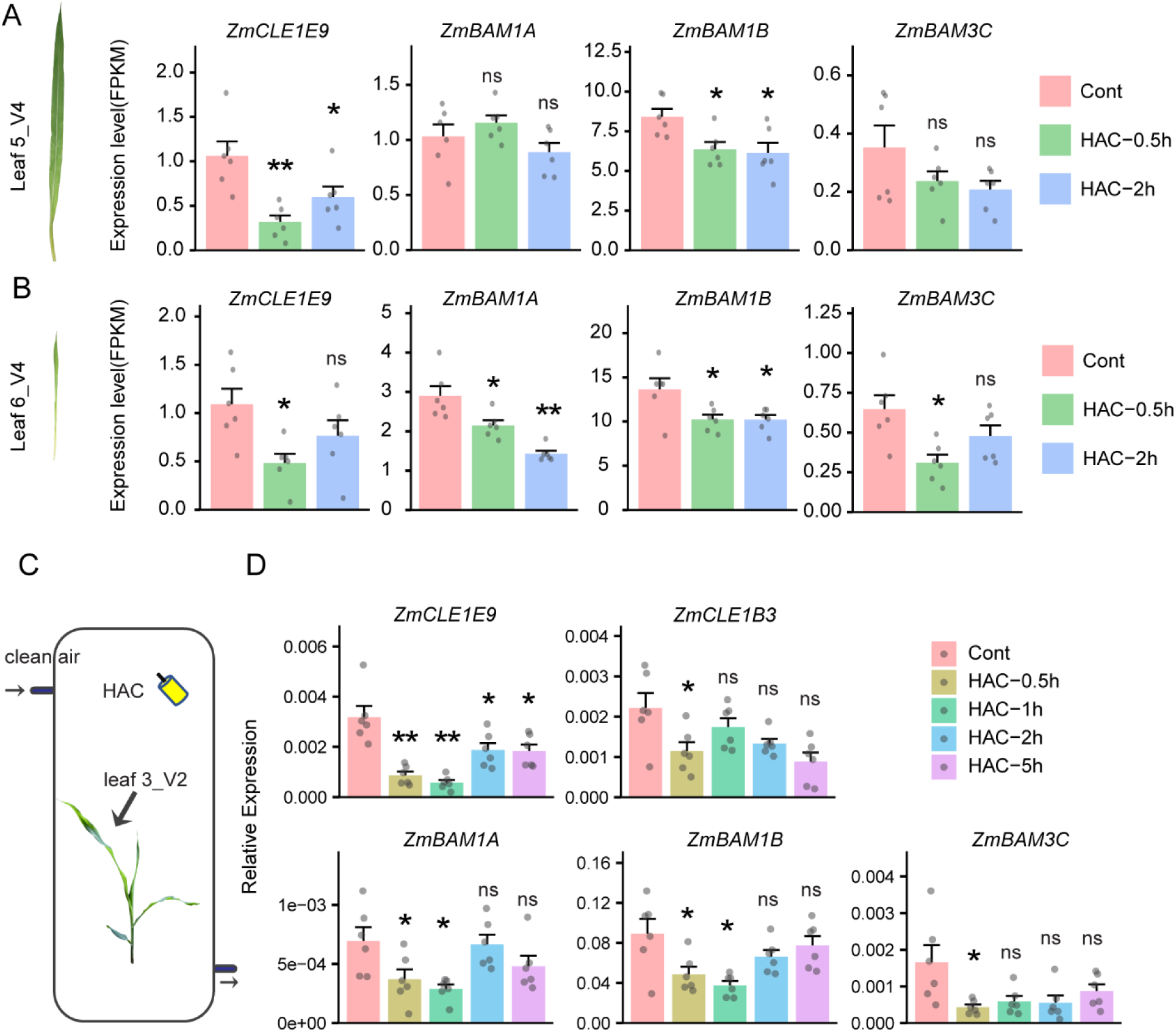
HAC exposure down-regulates CLE peptide signaling. (A) Gene expression level in HAC-exposed detached leaf 5 of V4 maize plants. (B) Gene expression level in detached leaf 6 of V4 maize plants exposed to HAC. (C) Experimental setup of V2 maize seedlings exposed to HAC; (D) Relative gene expression level in leaf 3 of HAC-exposed V2 maize seedlings. Data in A, B and D are presented as mean + s.e.(n = 6). Asterisks indicate significant difference compared to control (Cont) treatment: * P<0.05, ** P<0.01, two-tailed, two sample t test; ns: not significant. Data in A and B are retrieved from published data sets (Wang *et al*, 2023). Data in D are representative of 3 independent experiments with similar results.

**Figure EV1.**
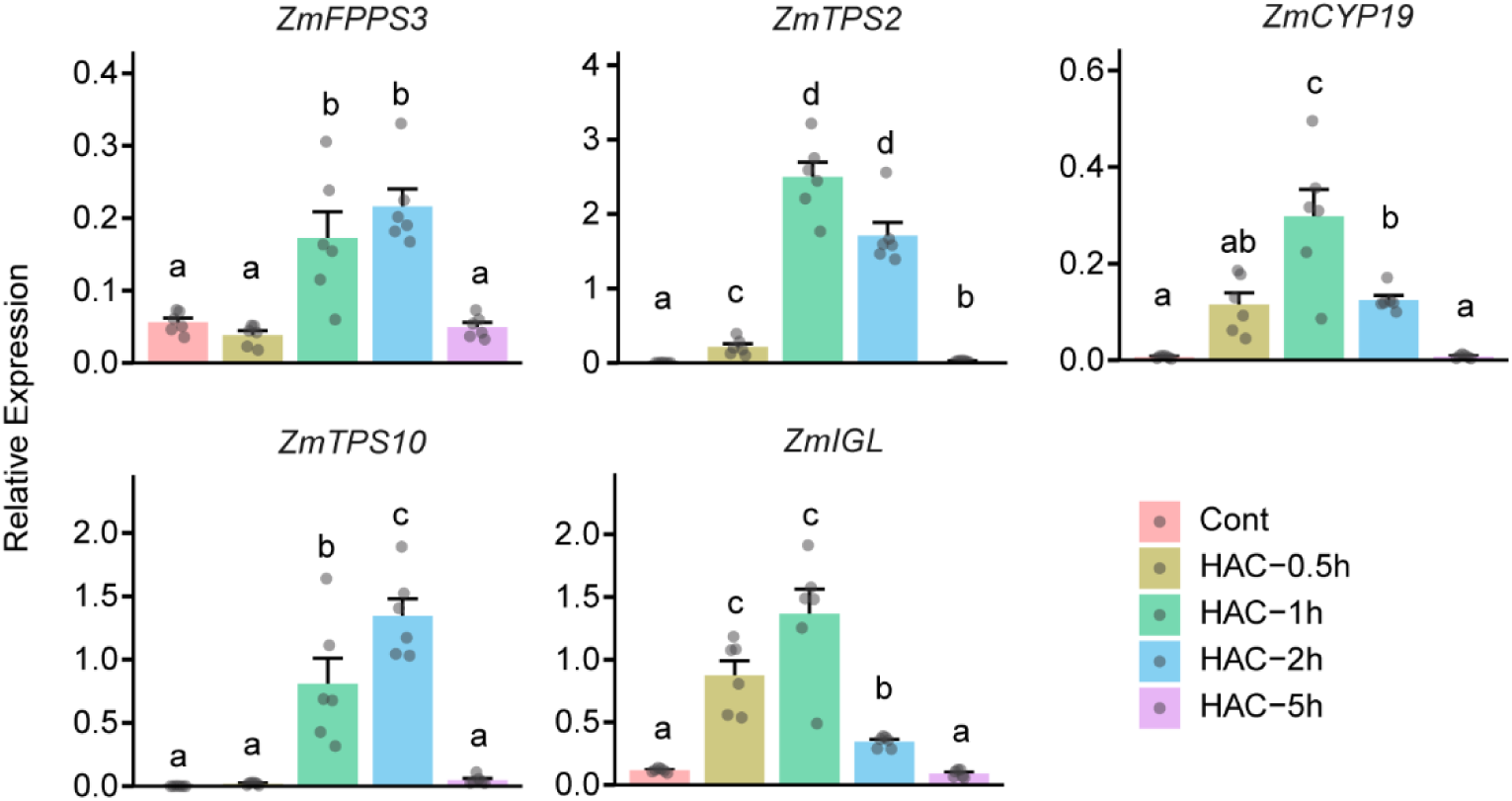
Relative expression of volatile biosynthesis genes in a HAC exposure time series experiment. Different letters above the bar indicate statistical differences. Differences in gene expression over time was determined using one-way ANOVA, followed with multiple comparisons tests (Tukey adjusted). Differences in *ZmIGL* expression were determined using a Kruskal-Wallis test. Data are shown as mean + s.e. (n = 6).

### Exogenous ZmCLE1E9 suppresses HAC triggered volatile emissions

To test if the CLE peptides regulate HAC-induced volatile emissions, we fed detached V2 stage B73 maize plants with peptide solutions and exposed them to HAC (Figure 2A). Compared to control plants, ZmCLE1E9-treated plants emitted significantly less indole, monoterpenes, sesquiterpenes and the homoterpenes DMNT and TMTT (Figure 2B). To gain further insights into ZmCLE1E9’s function, we generated *Zmcle1e9* mutant through CRISPR-Cas9 gene editing. In a pooled mutagenesis population based on the Hi-II background, we identified a mutant with 2 bp deletion in the coding sequence of the ZmCLE1E9 precursor gene (Figure EV2A). Because Hi-II is a hybrid between B73 and A188, we backcrossed this mutant to B73 for two generations before generating BC2F2. We then selected two homozygous mutant lines and a wild-type (WT) line from the segregation population to propagate seeds for further experiments (Figure EV2B). These mutants grew similarly to the WT plants (Figure 2C). When exposed to HAC, the *Zmcle1e9* mutants showed increased emissions of indole, monoterpenes, DMNT and TMTT compared to the WT plants (Figure 2D). When treated with simulated herbivory, only DMNT emissions were different (Figure EV2C). The peptide elicitor ZmPep3 induced volatile emissions in the mutants at similar levels as the WT plants (Figure EV2D). In contrast to ZmCLE1E9 pretreatment, ZmCLE1B3 pretreatment did not affect the emission of all the five groups of volatiles that we monitored (Figure EV3A). Additionally, HAC-induced volatile emissions did not increase in the *Zmcle1b3* mutants. Instead, HAC-induced emissions of monoterpenes and sesquiterpenes decreased in the *Zmcle1b3* plants (Figure EV3B-3D). Collectively, these results suggest that ZmCLE1E9 functions as a specific negative regulator of HAC-induced volatile emissions.

**Figure 2.**
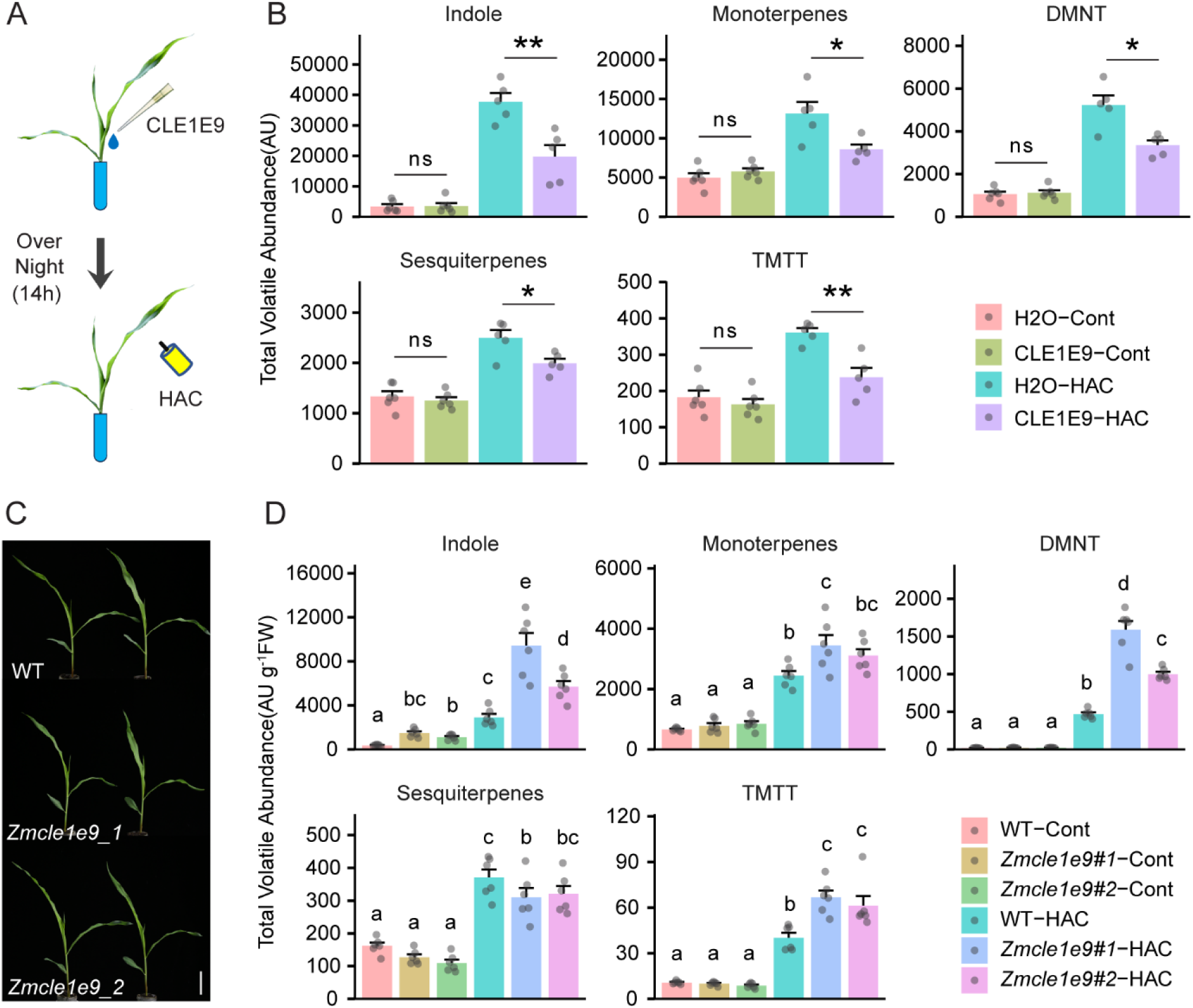
ZmCLE1E9 suppresses HAC-induced volatile emissions. (A) Schematic illustration of ZmCLE1E9 and HAC duo treatment. Detached V2 B73 seedlings were treated with 5 µM ZmCLE1E9, incubated overnight (14 h), before exposed to HAC dispensers. Water was used as control for ZmCLE1E9 treatment. No dispenser was used as control for HAC treatment. (B) Total volatile emissions of detached B73 seedlings with ZmCLE1E9 and HAC duo treatments. Relative emission rates of indole, monoterpenes, DMNT, sesquiterpenes and TMTT were determined by PTR-MS every half hour for 6 hours after HAC treatment and summed up as total volatile emissions. Asterisks indicate significant differences: * P<0.05, ** P<0.01, two-tailed, two sample t test; ns: not significant. AU: artificial unit. Data are shown as mean + s.e. (n = 5 or 6). (C) Growth phenotype of V2 stage *Zmcle1e9* mutants. Scale bar: 5cm. (D) Total volatile emissions of *Zmcle1e9* mutants after HAC exposure. Relative volatile emission rates were determined by PTR-MS every half hour for 6 hours after HAC treatment, normalized by plant fresh weight and then summed up as total volatile emissions. For each compound differences in emission between treatments and genotype were determined using two-way ANOVA (Type II). For monoterpenes differences were determined using heteroscedasticity-consistent standard errors obtained by White-adjusted ANOVAs (White, 1980). ANOVAs were conducted on log-transformed emission values for indole, DMNT and TMTT. Different letters represent significant differences as determined by multiple comparisons tests (Tukey adjusted). Data are shown as mean + s.e. (n = 6).

**Figure EV2.**
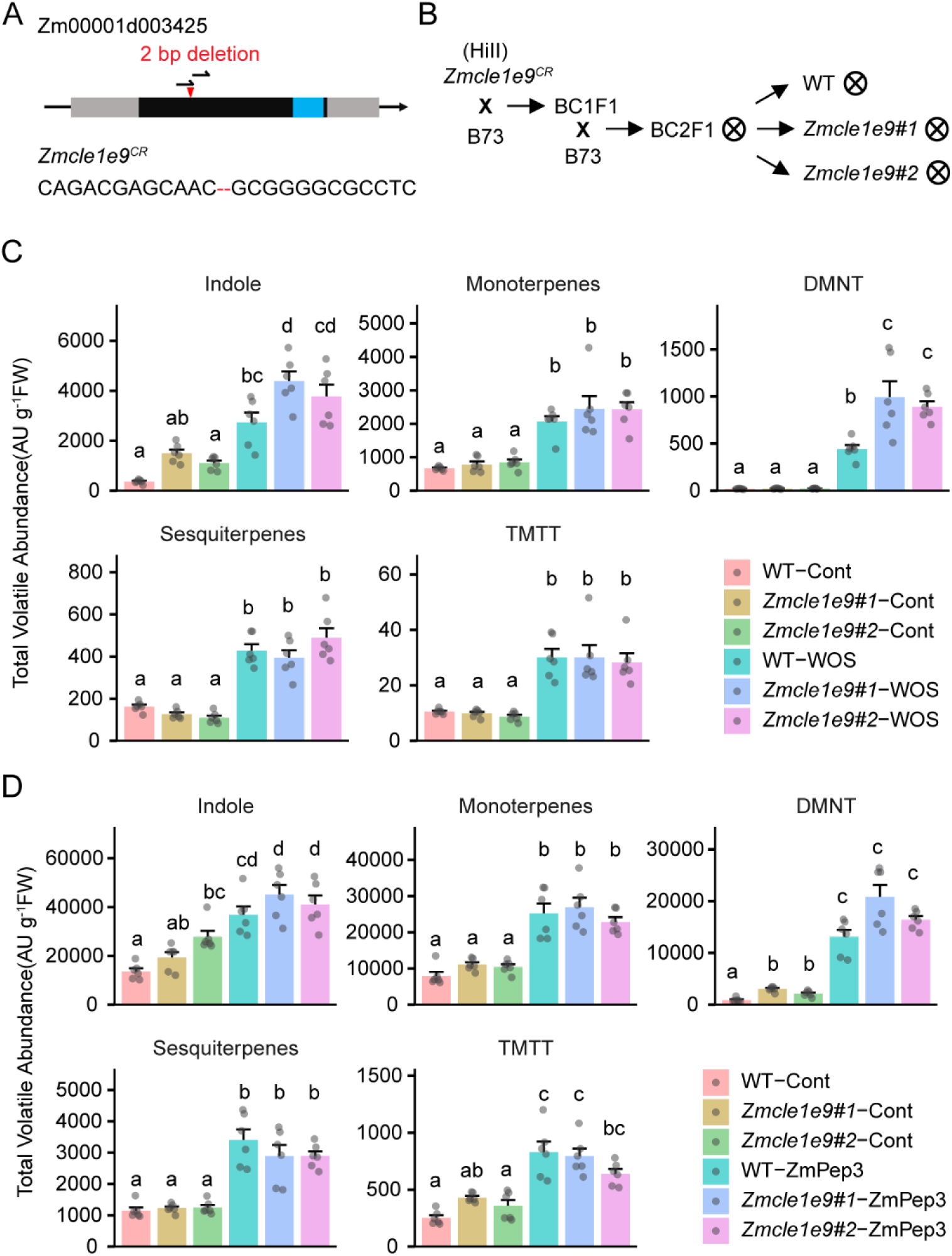
(A) Gene model, gRNA target sites (black arrows) and *ZmCLE1E9* CRISPR-Cas9 allele. Red triangle indicates the position of gene editing. Blue bar indicates the coding sequence for the predicted mature CLE1E9 peptide. (B) Genealogy of *Zmcle1e9* mutant. (C) Total volatile emissions of *Zmcle1e9* after simulated herbivory (wounding + oral secretion, WOS). Relative volatile emission rates were determined by PTR-MS every half hour for 6 hours after WOS treatment, normalized by plant fresh weight and then summed up as total volatile emissions. For each compound differences in emission between treatments and genotype were determined using two-way ANOVA (Type II). For indole, sesquiterpenes and DMNT, differences were determined using heteroscedasticity-consistent standard errors obtained by White-adjusted ANOVAs (White, 1980). ANOVAs were conducted on log-transformed emission values for monoterpenes, DMNT and TMTT. Different letters represent significant differences as determined by multiple comparisons tests (Tukey adjusted). Data are shown as mean + s.e. (n = 6). (D) Total volatile emissions of detached V3 stage *Zmcle1e9* plants after ZmPep3 (1 µM) treatment. Relative volatile emission rates were determined by PTR-MS every half hour for 6 hours after WOS treatment, normalized by plant fresh weight and then summed up as total volatile emissions. For each compound differences in emission between treatments and genotype were determined using two-way ANOVA (Type II). For monoterpenes and sesquiterpenes, differences were determined using heteroscedasticity-consistent standard errors obtained by White-adjusted ANOVAs (White, 1980). ANOVAs were conducted on log-transformed emission values for DMNT. Different letters represent significant differences as determined by multiple comparisons tests (Tukey adjusted). Data are shown as mean + s.e. (n = 6).

**Figure EV3.**
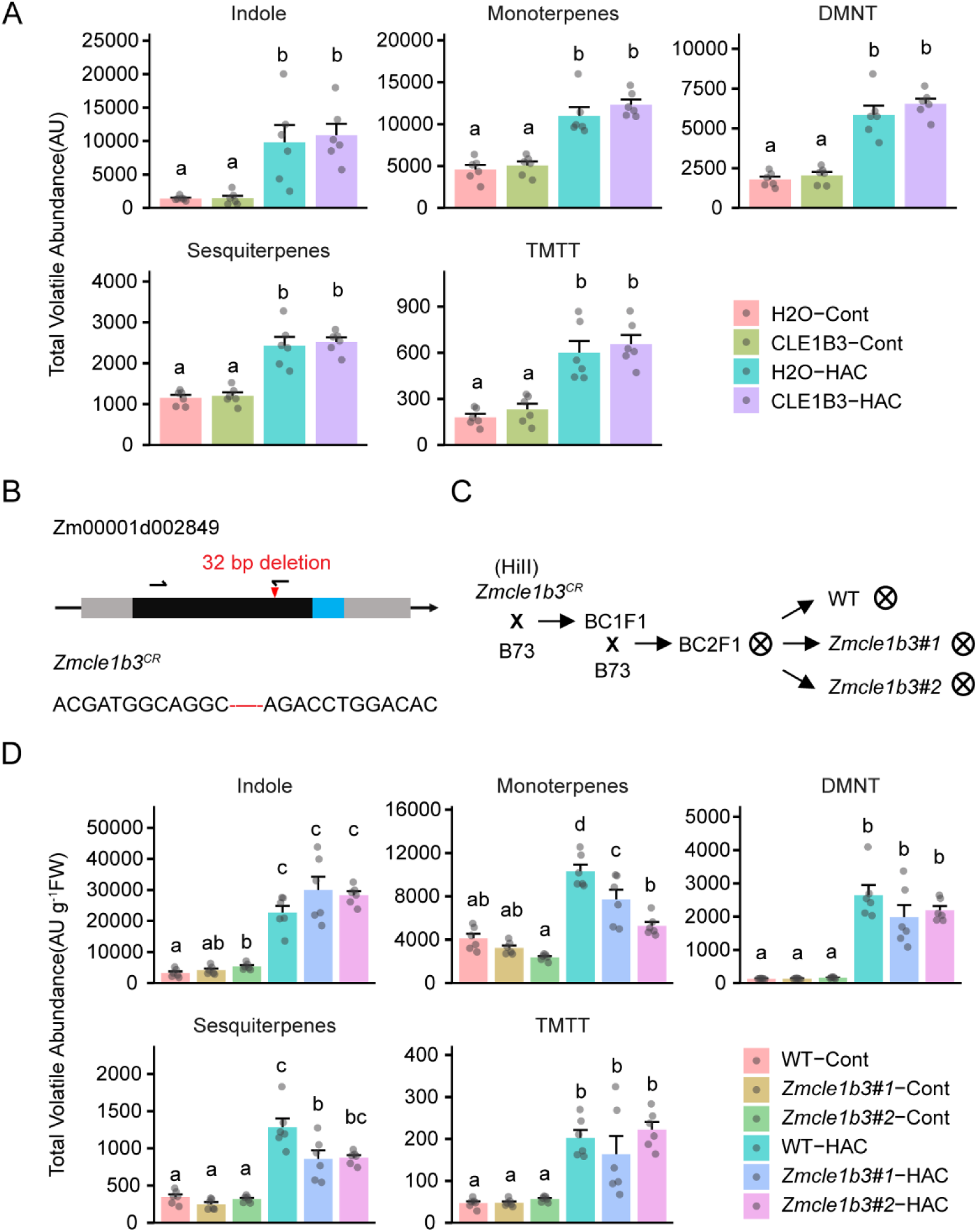
(A) Total volatile emissions of detached B73 seedlings with ZmCLE1B3 (5 µM) and HAC duo treatments. Relative emission rates of indole, monoterpenes, DMNT, sesquiterpenes and TMTT were determined by PTR-MS every half hour for 6 hours after HAC treatment and summed up as total volatile emissions. For each compound differences in emission between treatments and genotype were determined using two-way ANOVA (Type II). For TMTT, differences were determined using heteroscedasticity-consistent standard errors obtained by White-adjusted ANOVAs. ANOVAs were conducted on log-transformed emission values for indole, and monoterpenes. Different letters represent significant differences as determined by multiple comparisons tests (Tukey adjusted). Data are shown as mean + s.e. (n = 6). (B) Gene model, gRNA target sites (black arrows) and *ZmCLE1B3* CRISPR-Cas9 allele. Red triangle indicates the position of gene editing. Blue bar indicates the coding sequence for the predicated mature CLE1B3 peptide. (C) Genealogy of *Zmcle1b3* mutant. (D) Total volatile emissions of *Zmcle1b3* mutants after HAC exposure. Relative volatile emission rates were determined by PTR-MS every half hour for 6 hours after HAC treatment, normalized by plant fresh weight and then summed up as total volatile emissions. For each compound differences in emission between treatments and genotype were determined using two-way ANOVA (Type II). For monoterpenes, DMNT, sesquiterpenes and TMTT, differences were determined using heteroscedasticity-consistent standard errors obtained by White-adjusted ANOVAs. ANOVAs were conducted on log-transformed emission values for indole, DMNT, sesquiterpenes and TMTT. Different letters represent significant differences as determined by multiple comparisons tests (Tukey adjusted). Data are shown as mean + s.e. (n = 6).

### ZmCLE1E9 functions through BAM receptors to regulate volatile emissions

In *Arabidopsis*, BAM receptors have been shown to bind CLE peptides and transduce CLE-triggered cellular signaling (Zhang *et al*, 2016; Qian *et al*, 2018; Roman *et al*, 2022). In HAC exposed maize leaves, the transcriptional changes of *ZmBAM1A*, *ZmBAM1B* and *ZmBAM3C* were similar to that of *ZmCLE1E9* (Figure 1A). This prompted us to test if these BAM receptors function with ZmCLE1E9 during HAC-induced signaling. We first used molecular docking to test if these BAM receptors can form receptor-ligand pairs with ZmCLE1E9. Overall, ZmBAM1A, ZmBAM1B and ZmBAM3C share very conserved binding pockets with the *Arabidopsis* CLE peptide receptor AtPXY and AtBAM3(Figure 3A, 3B, Figure EV4). These binding pockets all contain a highly coordinated N-terminal region through hydrogen bonds and a small hydrophobic patch important for ZmCLE1E9 binding (Figure EV4). Compared to other CLE receptors, ZmBAM3C has a different leucine-rich-repeat domain at the end of the binding pocket with one conserved arginine missing. This feature may reduce its binding ability to ZmCLE1E9 (Figure EV4A).

**Figure 3.**
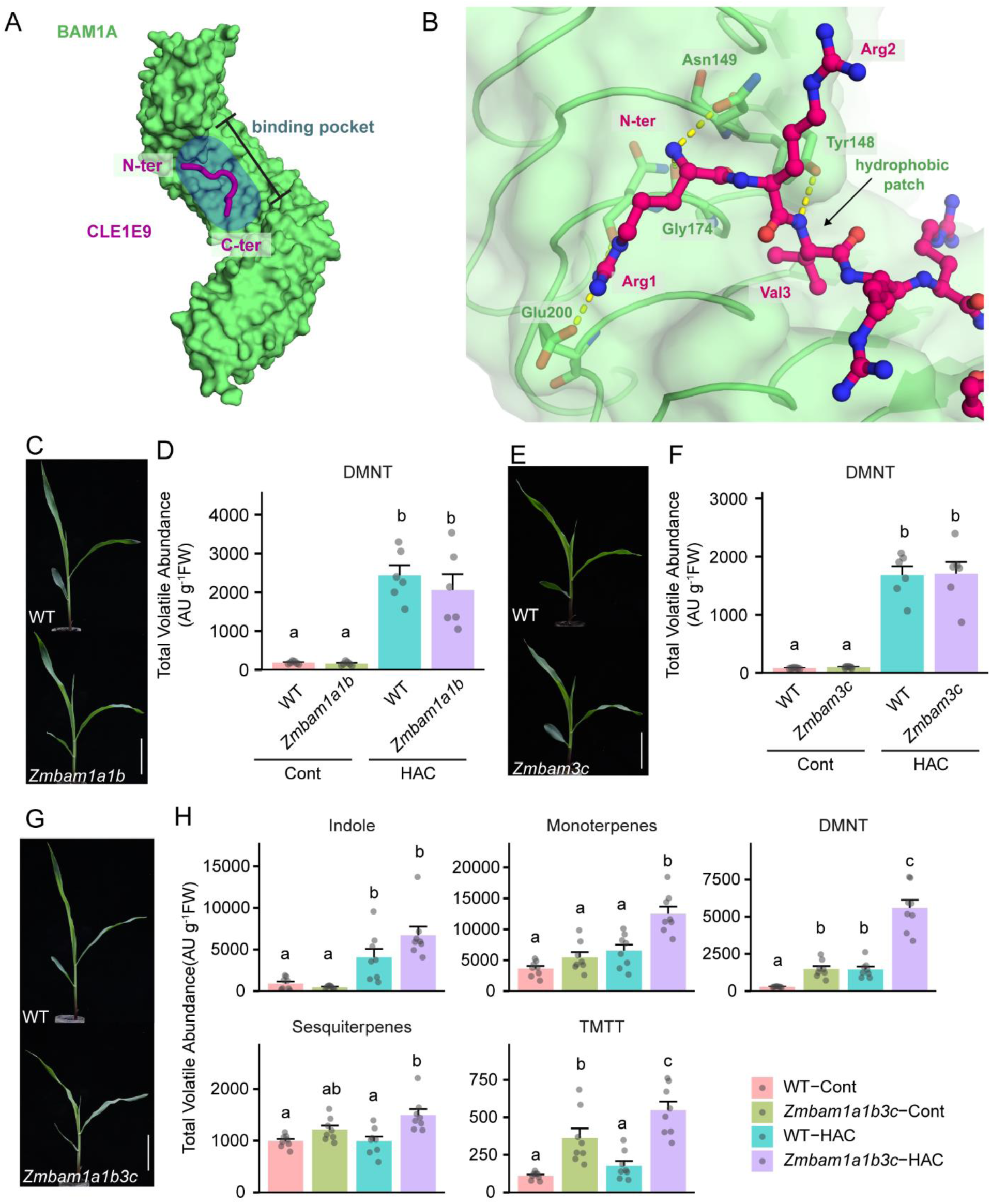
ZmBAM receptors form a molecular module with ZmCLE1E9 to regulate induced volatile emissions. (A) Overall view of the ZmCLE1E9-ZmBAM1A docking model. The ZmCLE1E9 peptide (shown in magenta) locates in the conserved receptor binding pocket (surface highlighted with a watermark in blue). The surface in green corresponds to the ectodomain of the ZmBAM1A model. (B) Close up view of the ZmCLE1E9-ZmBAM1A docking model. The N-terminal part of the ZmCLE1E9 peptide (shown in magenta) makes important contacts with the receptor (in green) and shares conserved features among other CLE-receptor pairs: a highly coordinated N-terminal region through hydrogen bonds and a small hydrophobic patch. The residue numbering corresponds to the position in the 12mer peptide. (C) Growth phenotype of V2 stage *Zmbam1a1b* double mutant. Scale bar: 5cm. (D) Total DMNT emission of *Zmbam1a1b* double mutant after HAC exposure. Relative volatile emission rates were determined by PTR-MS every half hour for 6 hours after HAC treatment, normalized by plant fresh weight and then summed up as total volatile emissions. Differences in DMNT emission between treatments and genotype were determined using two-way ANOVA (Type II). ANOVAs were conducted on log-transformed emission values. Different letters represent significant differences as determined by multiple comparisons tests (Tukey adjusted). Data are shown as mean + s.e. (n = 6). (E) Growth phenotype of V2 stage *Zmbam3c* double mutant. Scale bar: 5cm. (F) Total DMNT emission of *Zmbam3c* mutant after HAC exposure. Relative volatile emission rates were determined by PTR-MS every half hour for 6 hours after HAC treatment, normalized by plant fresh weight and then summed up as total volatile emissions. Differences in DMNT emission between treatments and genotype were determined using a Scheirer-Ray-Hare test. Different letters represent significant differences as determined by pairwise Wilcox tests. Data are shown as mean + s.e. (n = 6). (G) Growth phenotype of V2 stage *Zmbam1a1b3c* triple mutant. Scale bar: 5cm. (H) Total volatile emissions of *Zmbam1a1b3c* triple mutant after HAC exposure. Relative volatile emission rates were determined by PTR-MS every half hour for 6 hours after HAC treatment, normalized by plant fresh weight and then summed up as total volatile emissions. For each compound, differences in emission between treatments and genotype were determined using two-way ANOVA (Type II). For TMTT and indole, differences were determined using heteroscedasticity-consistent standard errors obtained by White-adjusted ANOVAs (White, 1980). ANOVAs were conducted on log-transformed emission values for indole and DMNT. Different letters represent significant differences as determined by multiple comparisons tests (Tukey adjusted). Data are shown as mean + s.e. (n = 8).

To understand the function of BAM receptors in HAC induced volatile emissions, we generated mutants using similar strategies as for the *Zmcle1e9* mutants. Because ZmBAM1A and ZmBAM1B are close homologs with 92% amino acid sequence identity, we generated a *Zmbam1a/Zmbam1b* (*Zmbam1a1b*) double mutant to test its ability to emit volatiles after HAC exposure (Figure EV4A, Figure EV5A-5C). The double mutant grew similarly to the WT plant. After HAC exposure, its volatile emissions were also similar as the WT plant. The emission of monoterpenes was lower than the WT plant (Figure 3C, 3D, Figure EV5D). Similarly, *Zmbam3c* did not show growth difference at seedling stage. HAC-induced volatile emissions were unchanged in the *Zmbam3c* single mutant (Figure 3E, 3F, Figure EV5E-5G). In *Arabidopsis*, several receptors function redundantly to recognize various CLE peptides (Zhang *et al*, 2016; Qian *et al*, 2018; Crook *et al*, 2020). Based on the volatile phenotype in the *Zmbam1a1b* double mutant and *Zmbam3c* single mutant, this is likely the case for the recognition of ZmCLE1E9 in maize. To test this hypothesis, we crossed *Zmbam1a1b* and *Zmbam3c* to generate *Zmbam1a/Zmbam1b/Zmbam3c* (*Zmbam1a1b3c*) triple mutant and proceeded to check its volatile phenotype. After HAC exposure, we observed significant increases in the emission of monoterpenes, DMNT, sesquiterpenes and TMTT in the triple mutant, similar to those observed in the *Zmcle1e9* single mutant. Interestingly, the emission of DMNT, sesquiterpenes and TMTT was increased in the triple mutant already before HAC exposure (Figure 3G, 3H). This may be caused by developmental differences in the triple mutant. The *Zmbam1a1b3c* mutant showed a more compacted architecture compared to its WT relative (Figure 3G). These results show that BAM receptors are negative regulators of HAC-induced volatile emissions in maize, possibly via CLE1E9 signaling.

**Figure EV4.**
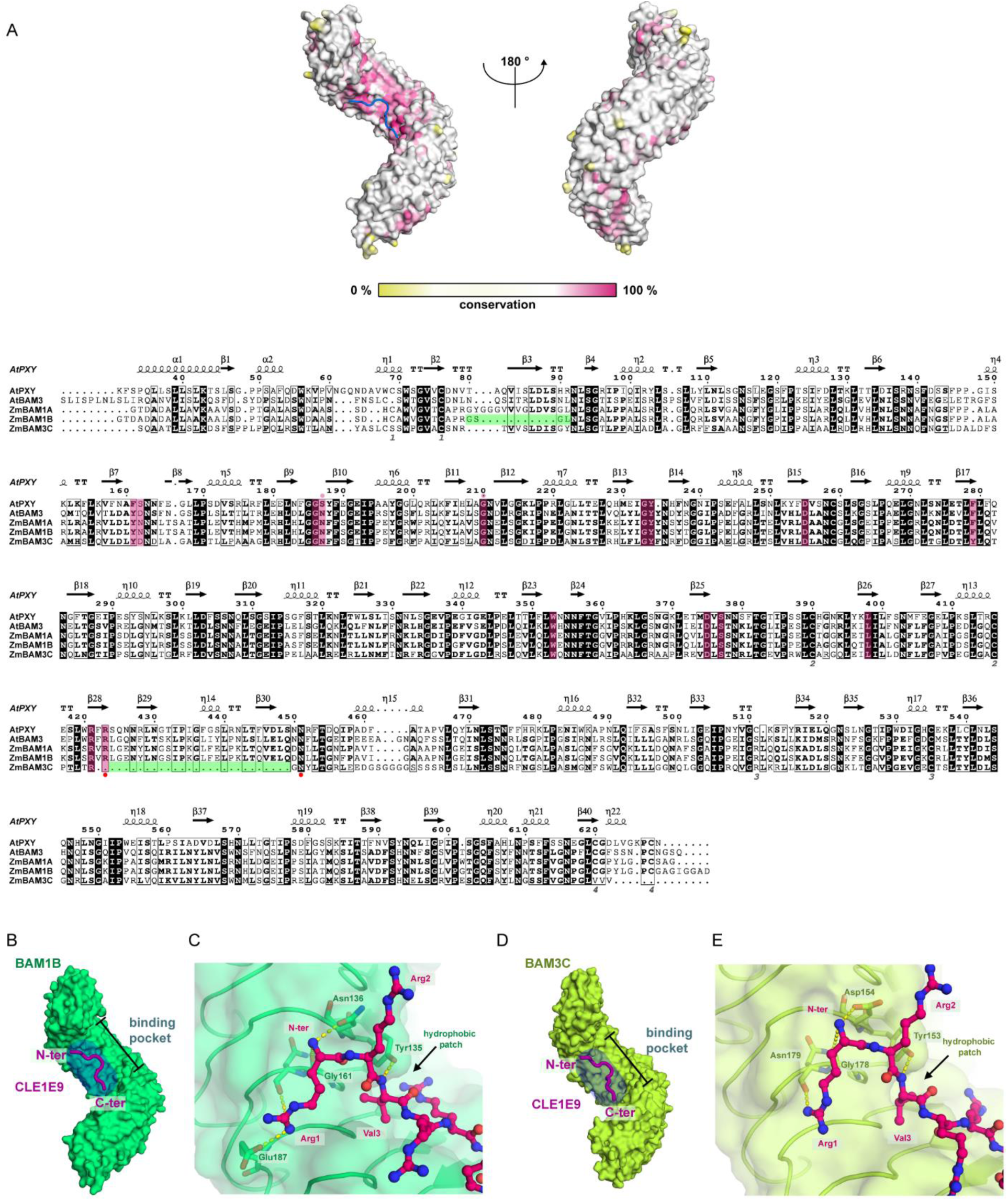
(A) Protein sequence alignment and conservation analysis of ZmBAM1A, ZmBAM1B and ZmBAM3C with the Arabidopsis receptors AtPXY and AtBAM3. The surface of the ectodomain of PXY is colored according to conservation among the 5 receptors. (B) Overall view of the ZmCLE1E9-ZmBAM1B docking model. (C) Close up view of the ZmCLE1E9-ZmBAM1B docking model. (D) Overall view of the ZmCLE1E9-ZmBA3C docking model. (E) Close up view of the ZmCLE1E9-ZmBAM3C docking model.

**Figure EV5.**
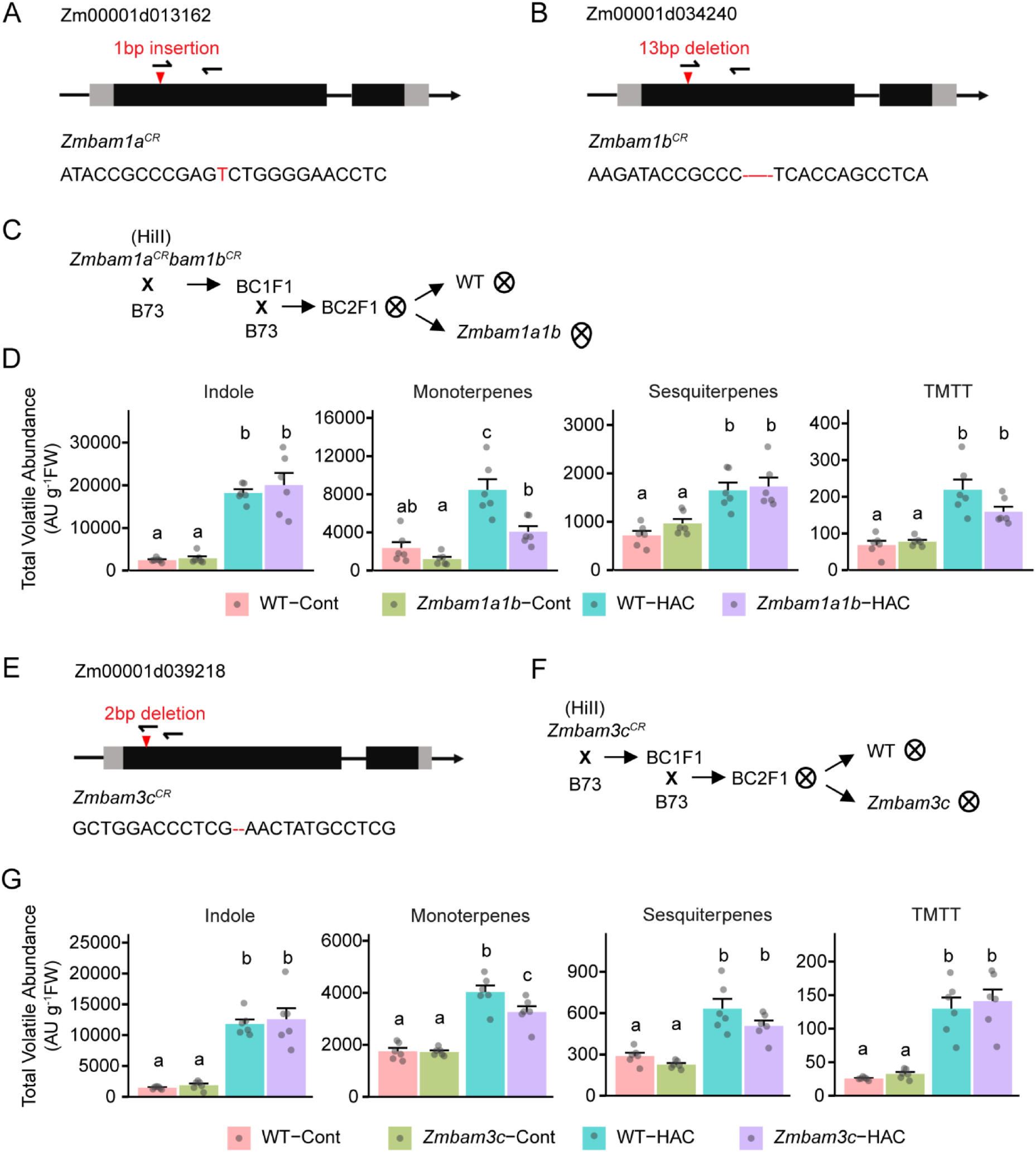
(A) Gene model, gRNA target sites (black arrows) and *ZmBAM1A* CRISPR-Cas9 allele. (B) Gene model, gRNA target sites (black arrows) and *ZmBAM1B* CRISPR-Cas9 allele. (C) Genealogy of *Zmbam1a1b* mutant. (D) Total volatile emissions of *Zmbam1a1b* mutants after HAC exposure. Relative volatile emission rates were determined by PTR-MS every half hour for 6 hours after HAC treatment, normalized by plant fresh weight and then summed up as total volatile emissions. Differences in emission between treatments and genotype were determined using two-way ANOVA (Type II). Log-transformed emission values were used for indole. Different letters represent significant differences as determined by multiple comparisons tests (Tukey adjusted). Data are shown as mean + s.e. (n = 6). (E) Gene model, gRNA target sites (black arrows) and *ZmBAM3C* CRISPR-Cas9 allele. (F) Genealogy of Zmbam3c mutant. (G) Total volatile emissions of Zmbam3c mutants after HAC exposure. Relative volatile emission rates were determined by PTR-MS every half hour for 6 hours after HAC treatment, normalized by plant fresh weight and then summed up as total volatile emissions. For each compound differences in emission between treatments and genotype were determined using two-way ANOVA (Type II). For sesquiterpenes, differences were determined using heteroscedasticity-consistent standard errors obtained by White-adjusted ANOVAs. ANOVAs were conducted on log-transformed emission values for indole and TMTT. Different letters represent significant differences as determined by multiple comparisons tests (Tukey adjusted). Data are shown as mean + s.e. (n = 6).

### The ZmCLE1E9-ZmBAMs module regulates stomatal aperture

Induced volatile emissions after HAC exposure are likely determined by many factors ranging from volatile biosynthesis to release via the stomata. We first tested if ZmCLE1E9 could affect HAC-induced volatile biosynthesis, using the expression of *ZmFPPS3*, *ZmTPS2*, *ZmCYP19*, *ZmTPS10 and ZmIGL* as well-established volatile biosynthesis genes whose expression is a rate-limiting step in volatile production (Seidl-Adams *et al*, 2015; Erb *et al*, 2015; Schnee *et al*, 2006; Richter *et al*, 2016). Pretreatment of ZmCLE1E9 did not affect the expression of all the genes tested, both before and after HAC exposure (Figure 4A, Figure EV6A). To confirm this pattern, we proceeded with checking the expression pattern of volatile biosynthesis genes in the mutants. In the *Zmcle1e9* mutants, all five genes showed similar or even lower expression level in the mutant compared to WT plants after HAC exposure. Gene expression patterns in the two mutant lines showed some variations (Figure EV6B). In the *Zmbam1a1b3c* triple mutant, only *ZmFPPS3* showed significantly higher expression after HAC exposure compared to WT (Figure EV6C). Together, these results suggest that the ZmCLE1E9-ZmBAM module is unlikely to regulate volatile release via transcriptional reprogramming of volatile biosynthesis. By inference, we can also exclude regulation of early HAC-induced defense signaling by the ZmCLE1E9-ZmBAM module, as this would be visible at the level of the expression of the tested biosynthesis genes (Hu *et al*, 2019).

**Figure 4.**
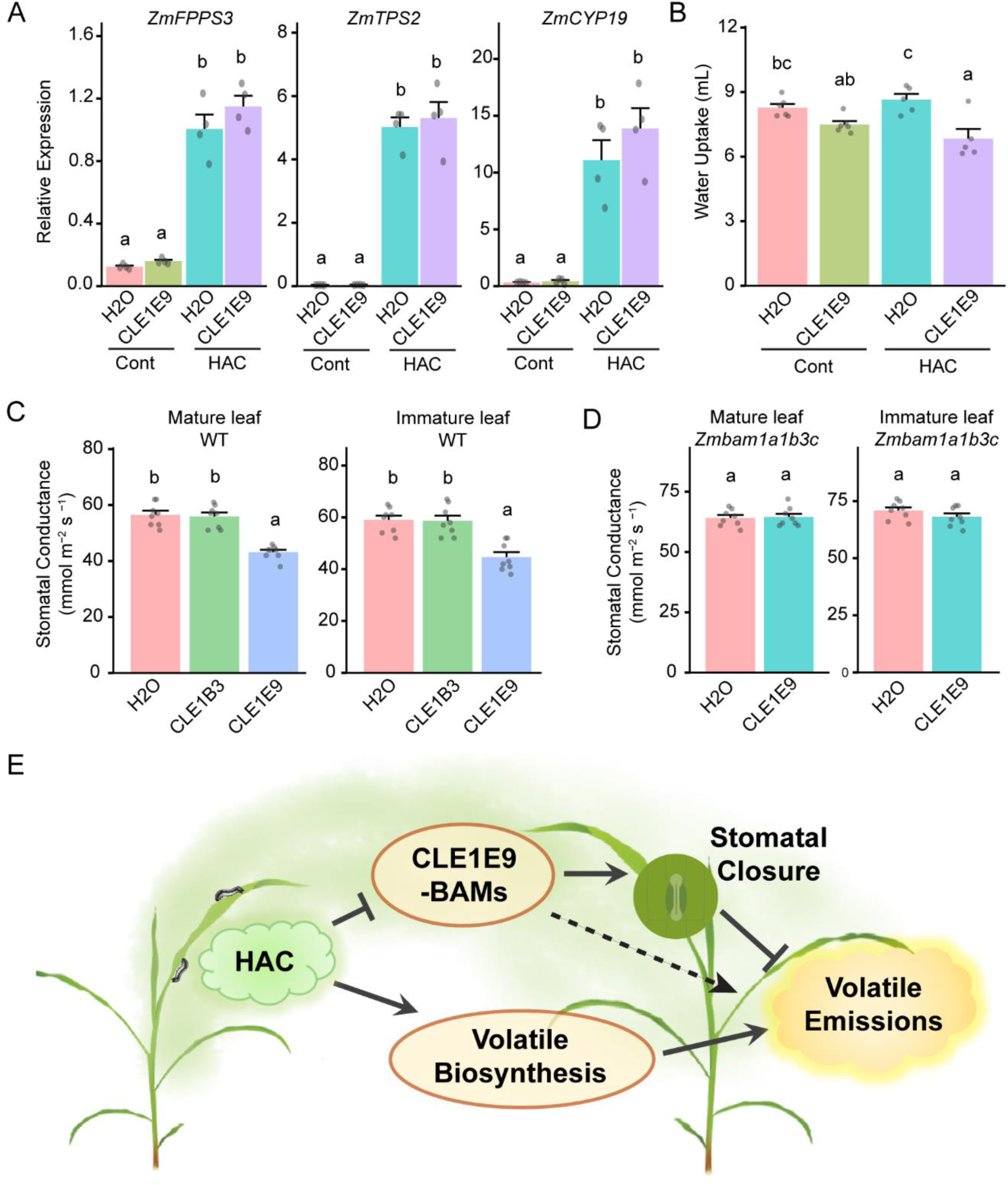
The CLE1E9-BAMs module suppresses volatile emissions by promoting stomatal closure. (A) Relative expression level of volatile biosynthesis genes in leaf 3 of V2 stage B73 maize plants treated with ZmCLE1E9 (5 µM) and HAC dispenser. Different letters above the bar indicate statistical differences. For each gene, differences in gene expression between treatments and genotype were determined using two-way ANOVA (Type II). ANOVAs were conducted on log-transformed emission values for *ZmTPS2* and *ZmCYP19*. Different letters represent significant differences as determined by multiple comparisons tests (Tukey adjusted). Data are shown as mean + s.e. (n = 4). (B) Water uptake of detached V2 stage B73 seedlings with ZmCLE1E9 and HAC treatment. Differences in water uptake between treatments and genotype were determined using a Scheirer-Ray-Hare test. Data are shown as mean + s.e. (n = 6). (C) Stomatal conductance of detached leaves from V4 stage B73 plants. Stomatal conductance of mature leaves (leaf 4) and immature leaves (leaf 5) was measured 2 hours after treatment with 5 µM peptide. Differences in stomatal conductance between treatments were determined using one-way ANOVA. Different letters represent significant differences as determined by multiple comparisons tests (Tukey adjusted). Data are shown as mean + s.e. (n = 8). (D) Stomatal conductance of detached leaves from V5 stage *Zmbam1a1b3c* plants. Stomatal conductance of mature leaves (leaf 5) and immature leaves (leaf 6) was measured 2 hours after treatment with 5 µM peptide. Differences in stomatal conductance were determined using Welch’s t-tests. Different letters above the bar indicate statistical differences. Data are shown as mean + s.e. (n = 8). (E) Schematic model of how the ZmCLE1E9-ZmBAMs module regulates HAC-induced volatile emissions. ZmCLE1E9 functions through the BAM receptors to close stomata. HAC exposure leads to reduced expression of the ZmCLE1E9-ZmBAMs molecular module. Together with the increased volatile biosynthesis, this regulation pattern contributes to increased volatile emissions. The dashed line indicates other possible mechanisms of the ZmCLE1E9-ZmBAMs in regulating HAC-induced volatile emissions.

GLV and HAC-induced volatile emissions are sensitive to stomatal aperture (Figure EV8) (Seidl-Adams *et al*, 2015). Thus, we explored the potential role of stomata in ZmCLE1E9-mediated volatile emissions. First, we tested if the increased volatile emissions after HAC exposure in the mutants is associated with altered stomatal development. Maize stomata are formed by two central dumbbell-shaped guard cells and two lateral subsidiary cells (Nunes *et al*, 2020). We did not observe major differences in stomatal morphology between the *Zmcle1e9* and *Zmbam1a1b3c* mutants and their respective WT siblings (Figure EV7). The stomata in the *Zmbam1a1b3c* mutant are less wide (Figure 7G). Stomatal density in the mutants was similar as in WT plants (Figure EV7E, 7H). In *Arabidopsis*, the peptide AtCLE9 and AtCLE25 induce stomatal closure (Takahashi *et al*, 2018; Zhang *et al*, 2019). Thus, we tested if ZmCLE1E9 regulates stomatal aperture in maize. As a first proxy for stomatal aperture and transpiration, we measured water uptake in detached leaves. ZmCLE1E9 feeding reduced water uptake of maize seedlings, with the strongest differences observed after HAC treatment (Figure 4B). We then measured stomatal conductance of maize leaves 2 hours after peptide feeding. In both mature and immature leaves of B73 (WT) leaves, ZmCLE1E9, but not ZmCLE1B3, significantly reduced stomatal conductance (Figure 4C). This effect disappeared in the *Zmbam1a1b3c* mutants (Figure 4D), confirming that the ZmCLE1E9 function depends on the ZmBAM receptors. Collectively, these data suggest that the ZmCLE1E9-ZmBAMs module acts as a negative regulator of stomatal conductance and thereby reduce volatile emission.

**Figure EV6.**
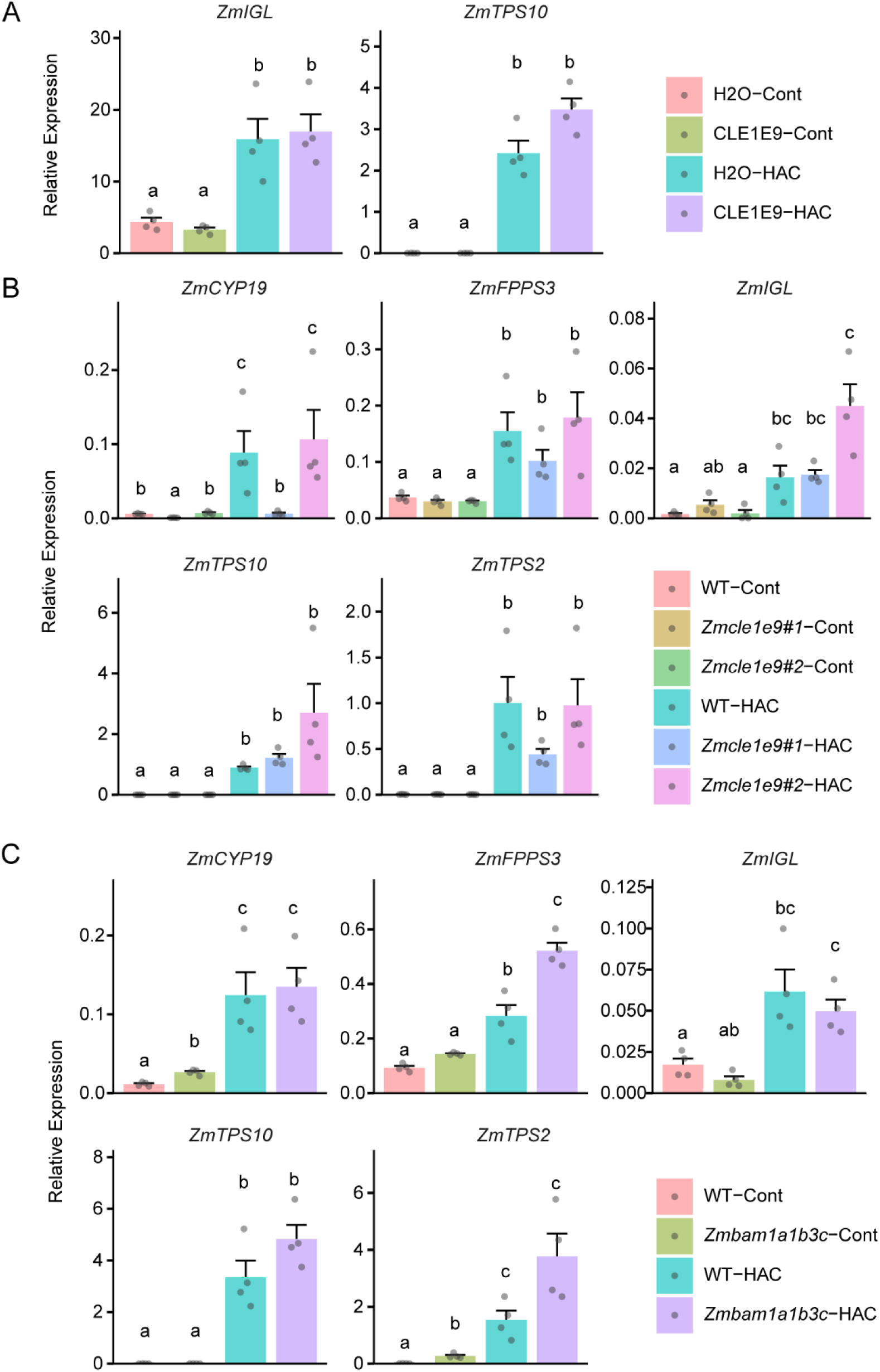
(A) Relative expression level of volatile biosynthesis genes in leaf 3 of V2 stage B73 maize plants treated with ZmCLE1E9 (5 µM) and HAC dispenser. Differences in *ZmIGL* expression between treatments and genotypes were determined using two-way ANOVA (type II). Differences in ZmTPS10 expression were determined using a Scheirer-Ray-Hare test. Different letters represent significant differences as determined by multiple comparisons tests (Tukey adjusted) and a pairwise Wilcox test for ZmIGL and ZmTPS10, respectively. Data are shown as mean + s.e. (n = 4). (B) Relative expression level of volatile biosynthesis genes in leaf 3 of HAC-exposed V2 stage *Zmcle1e9* mutants. For each gene, differences in gene expression between treatments and genotypes were determined using two-way ANOVA (type II). For *ZmIGL*, differences were determined using heteroscedasticity-consistent standard errors obtained by White-adjusted ANOVAs. ANOVAs were conducted on log-transformed emission values for *ZmCYP19*, *ZMFPPS3*, *ZmIGL* and *ZmTPS2*. Differences in *ZmTPS10* expression were determined using a Scheirer-Ray-Hare test. Different letters represent significant differences as determined by multiple comparisons tests (Tukey adjusted) for all genes except *ZmTPS10*, in which case a pairwise Wilcox test was used. Data are shown as mean + s.e. (n = 4). (C) Relative expression level of volatile biosynthesis genes in leaf 3 of HAC-exposed V2 stage *Zmbam1a1b3c* mutants. For each gene differences in gene expression between treatments and genotype were determined using two-way ANOVA (Type II). ANOVAs were conducted on log-transformed emission values for *ZmTPS2*, *ZmTPS10* and *ZmCYP19*. Different letters represent significant differences as determined by multiple comparisons tests (Tukey adjusted). Data are shown as mean + s.e. (n = 4).

**Figure EV7.**
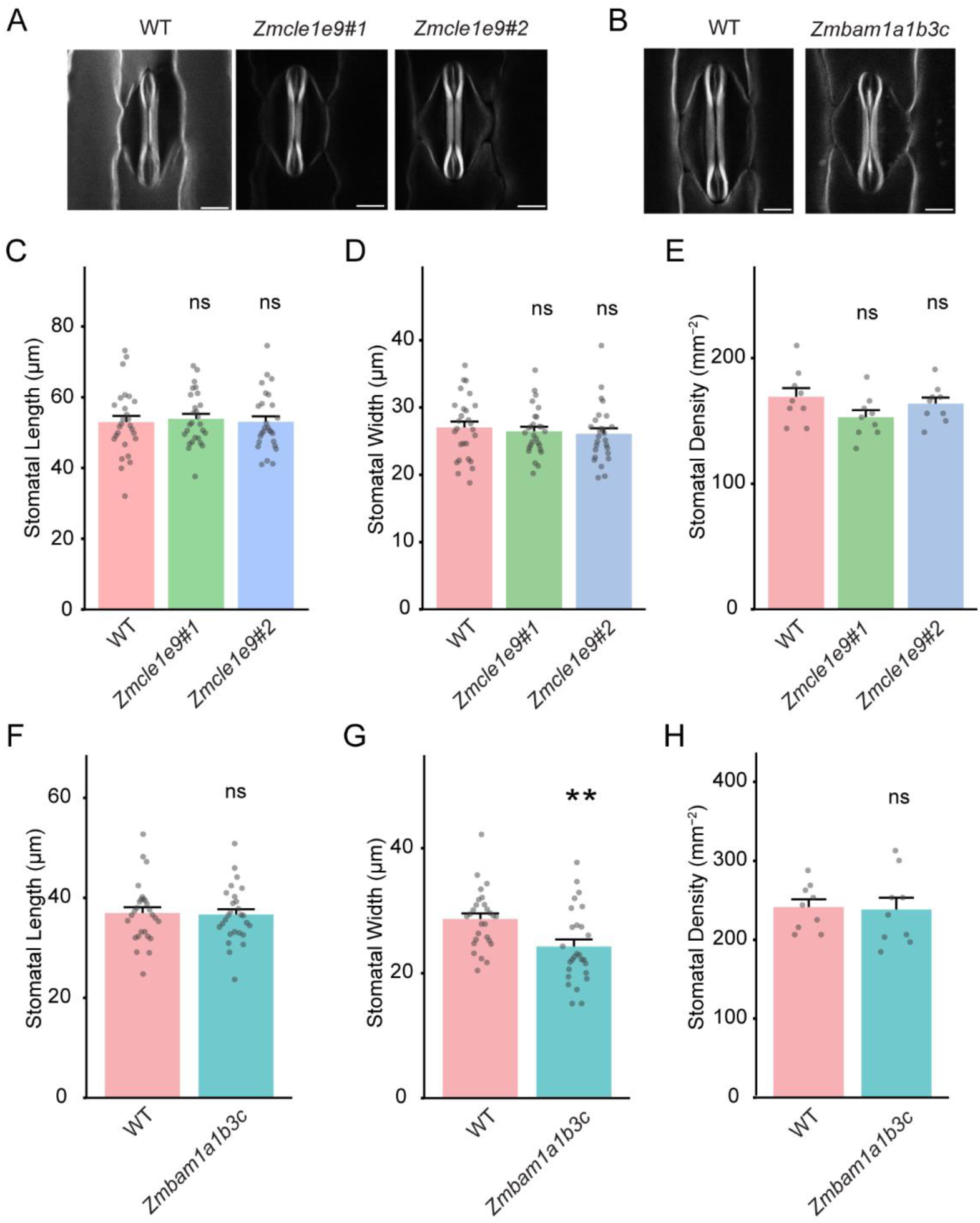
(A) XY sections of 3D imaged mature stomatal complexes in the *Zmcle1e9* mutants. Images were acquired from the second leaves of V2 stage plants. Scale bar: 10 µm. (B) XY sections of 3D imaged mature stomatal complexes in the *Zmbam1a1b3c* mutants. Images were acquired from the second leaves of V2 stage plants. Scale bar: 10 µm. (C)Length of stomatal complex from the second leaves of V2 stage *Zmcle1e9* mutants and the corresponding WT plants. Data are shown as mean + s.e. (n = 27). (D)Width of stomatal complex from the second leaves of V2 stage *Zmcle1e9* mutants and corresponding WT plants. Data are shown as mean + s.e. (n = 27). (E) Stomatal density of *Zmcle1e9* mutants. Stomatal density was calculated from the abaxial surface of leaf 2 of V2 plants. Data are shown as mean + s.e. (n = 9). (F) Length of stomatal complex from the second leaves of V2 stage *Zmbam1a1b3c* mutants and the corresponding WT plants. Data are shown as mean + s.e. (n = 27). (G) Width of stomatal complex from the second leaves of V2 stage *Zmbam1a1b3c* mutants and the corresponding WT plants. Data are shown as mean + s.e. (n = 27). (H) Stomatal density of *Zmbam1a1b3c* mutants. Stomatal density was calculated from the abaxial surface of leaf 2 of V2 plants. Differences in stomatal length, width and density were determined using one-way ANOVA, ns indicated no significant difference compared to the WT, double asterisks indicate significant difference compared to WT (** P<0.01, two-tailed, two sample t test). Data are shown as mean + s.e. (n = 9).

**Figure EV8.**
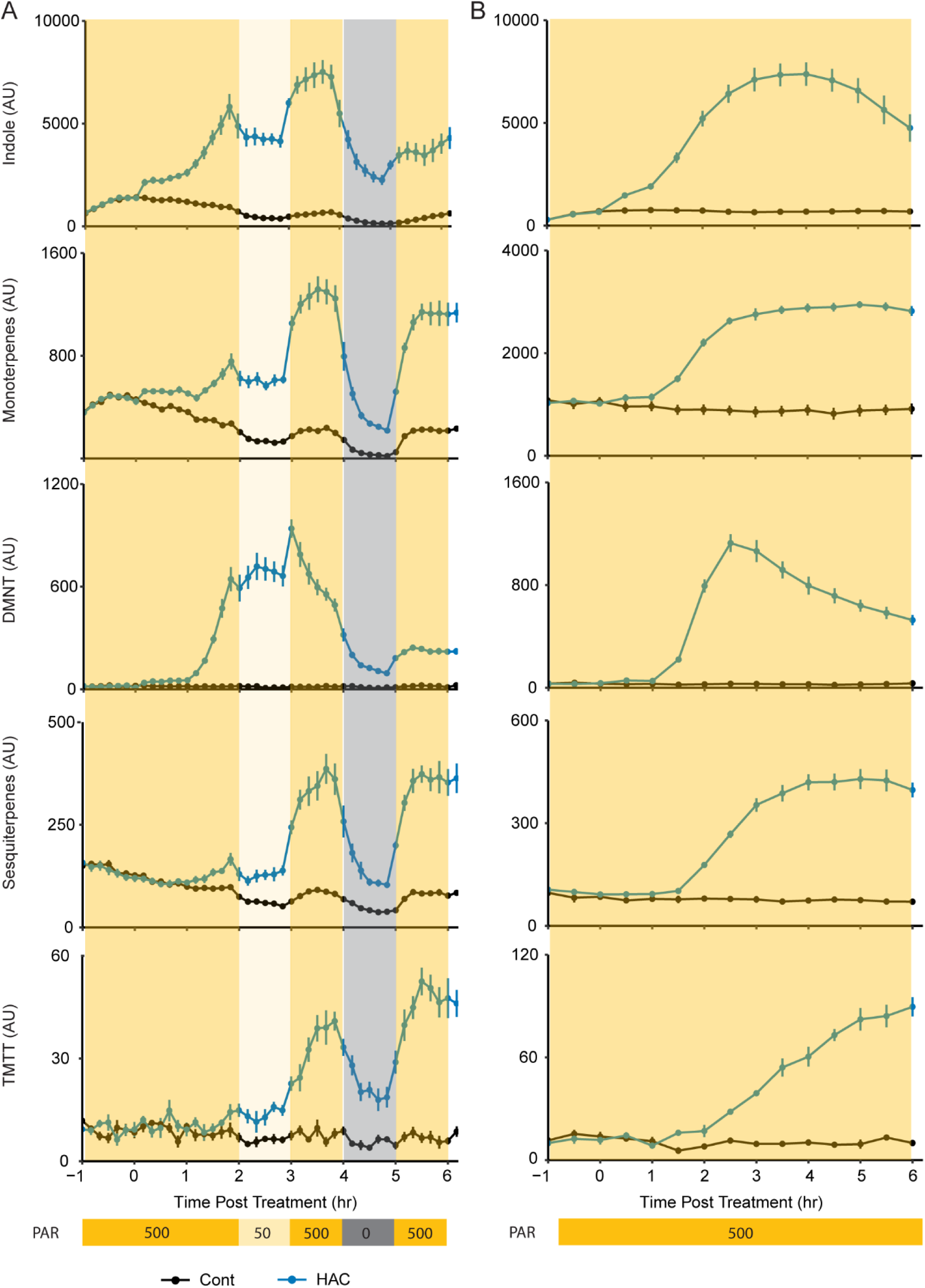
(A) Volatile emission kinetic of HAC-exposed V2 stage B73 seedling under dynamic light. Relative volatile emission rates were determined by PTR-MS every 10 minutes. Light scheme is shown at the bottom of the figure. AU: artificial unit. PAR: photosynthetically active radiation. Data are shown as mean ± s.e. (n = 6). (B) Volatile emission kinetic of HAC-exposed V2 stage B73 seedling under constant light. Relative volatile emission rates were determined by PTR-MS every 30 minutes. Light scheme is shown at the bottom of the figure. Data are shown as mean ± s.e. (n = 6).

## Discussion

In response to herbivore attack, plants engage with each other by emitting and perceiving volatile compounds (Turlings & Erb, 2018; Takabayashi & Shiojiri, 2019). Maize plants perceive herbivory induced volatiles such as HAC and respond by releasing terpenoids and indole (Engelberth *et al*, 2004; Wang *et al*, 2023). If and how volatile emissions are regulated in plant-plant interactions beyond canonical hormonal signaling and biosynthesis is not well understood. Here we show that the ZmCLE1E9 peptide functions through the ZmBAM receptors to suppress volatile emissions, most likely by reducing their release through the stomata. Upon HAC perception, receiver plants downregulate the ZmCLE1E9-ZmBAMs module, which allows them to release more volatiles themselves, thereby effectively enhancing population level volatile concentrations. Here, we discuss the molecular mechanisms and agroecological consequences of this phenomenon.

The CLE peptide family contains members involved in various aspects of plant development and responses to environmental stimuli. Their functions are often dependent on functionally redundant receptors. For instance, AtCLE25 needs both AtBAM1 and AtBAM3 to trigger stomatal closure (Takahashi *et al*, 2018). AtBAM1, AtBAM2 and AtBAM3 together regulates AtCLE16-mediated cell division in roots (Crook *et al*, 2020). In this work, we show that the activity of ZmCLE1E9 in regulating stomatal closure also requires three BAM receptors (Figure 3). The functions of the CLE-BAM signaling module largely depend on the spatial temporal expression patterns, as illustrated by the multifaceted AtBAM1 and AtBAM3 mentioned above (Takahashi *et al*, 2018; Crook *et al*, 2020). We found that the *Zmbam1a1b3c* showed altered seedling growth, smaller stomata and increased basal level of DMNT and TMTT emissions (Figure 2G and 2H, Figure EV7). It’s possible that next to regulating stomatal activity, BAM receptors control other developmental programs with a yet unclear link to volatile emissions in maize (Figure 4E). Conducting a comprehensive transcriptomic analysis at high spatial temporal resolution with the *bam* and *cle* mutants would shed light on the link between maize leaf development and volatile emissions.

The role of the ZmCLE1E9-ZmBAMs module in regulating volatile emissions appears to be specific to HAC perception, since simulated herbivory and ZmPep3-induced volatile emissions are largely unaffected in the *Zmcle1e9* mutants (Figure 2, Figure EV2B, 2C). This is in line with the notion that different herbivory-associated cues trigger distinct signaling pathways to fine-tune physiological responses (Erb & Reymond, 2019). Simulated herbivory and ZmPep3 are directly related to damage caused by herbivory (Waterman *et al*, 2019; Huffaker *et al*, 2013). In this scenario, water loss and potential pathogen entry through the damaged sites pose an imminent threat. A typical plant response against such a threat is stomatal closure. For example, *Nicotiana attenuata* plants close stomata upon attack by *Manduca sexta* larvae (Nabity *et al*, 2013; Meza-Canales *et al*, 2017). In *Arabidopsis*, the peptide AtPep1, a ZmPep3 relative, can trigger stomatal closure (Zheng *et al*, 2018). The dilemma to close stomata to counteract pest infection and to open stomata for emitting volatiles is then likely solved by recruiting other molecular modules. The *Arabidopsis* SCREWs-NUT module has been reported to reopen stomata in plant immunity (Liu *et al*, 2022). Homologs of SCREWs and NUT also exist in maize (Liu *et al*, 2022), whether they regulate volatile emissions triggered by herbivory remains to be tested. Volatile receiver plants do not yet suffer from mechanical damage caused by herbivory, and can thus keep their stomata open, which again maximizes their capacity to release induced volatiles.

It is now well accepted that plant volatiles play a central role in shaping tritrophic interactions and holds great potential to improve the efficiency of biological control in agriculture (Turlings & Erb, 2018; Takabayashi & Shiojiri, 2019). Attempts have been made to engineer crops with higher volatile emissions for increased herbivore resistance. For instance, (*E*)-β-caryophyllene emitted from insect-damaged maize is a major attractant of natural enemies (Rasmann *et al*, 2005; Köllner *et al*, 2008). Its biosynthesis gene *ZmTPS23* contributes to indirect defense in maize (Köllner *et al*, 2008). Overexpressing this gene in maize leads to constitutive emission of (*E*)-β-caryophyllene and α-humulene (Robert *et al*, 2013). However, this change increased plant apparency to the herbivore *Spodoptera fruigiperda* and led to more leaf damage in the field. In addition, the transgenic maize showed growth penalties including comprised germination rate and lower yield (Robert *et al*, 2013). Thus, exploiting the potential of volatiles in biological control requires more than simply producing more volatiles. Our results here show that maize plants have evolved sophisticated mechanisms to control stomata-governed volatile emissions. Previous work on petunia also highlighted the importance of an ABC transporter in volatile emissions (Adebesin *et al*, 2017). Rapid advancement in gene editing, protein engineering and volatile analysis together with a detailed understanding of the regulation of volatile release will greatly promote the understanding of plant volatile biology and pave the way for developing plants that release bioactive volatiles in the right place at the right time.

## Methods

| Reagents and tools table |  |  |
| --- | --- | --- |
| Reagent/resource | Reference or source | Identifier or catalog number |
| **Oligonucleotides and other sequence-based reagents** |  |  |
| Primers | This study | Table EV1 |
| Peptides | GeneScript | Table EV2 |
| **Chemicals, enzymes and other reagents** |  |  |
| (Z)-3-hexenyl acetate | Sigma-Aldrich | W317101 |
| GeneJET PCR Purification Kit | Fisher Scientific | K0702 |
| DNAse I | Fisher Scientific | EN0521 |
| PrimeScript™ RT Master Mix (Perfect Real Time) | TaKaRa | RR036A |
| ORA™ SEE qPCR Green ROX L Mix | AxonLab | 11006014 |
| Acetic acid | Sigma-Aldrich | 45754 |
| Direct Red23 | Sigma-Aldrich | 212490 |
| Ethanol | Sigma-Aldrich | E7023 |
| PBS | Sigma-Aldrich | P3813 |
| **Software** |  |  |
| Rstudio | <u>RStudio Desktop - Posit</u> |  |
| Fiji | (Schindelin *et al*, 2012) |  |
| PyMOL | https://pymol.org/ |  |

### Plant materials and growth conditions

Maize inbred line B73 was used in this study. The *Zmcle1e9*, *Zmcle1b3*, *Zmbam1a*, *Zmbam1b* and *Zmbam3c* mutants were generated as described below. All seeds were sown directly in commercial soil (Selmaterra, BiglerSamen, Switzerland) with a depth of ∼ 1 cm. Plants used at V2 stage were grown in plastic tubes (Ø 44 mm × H 114 mm). Plants used at V4 or V5 stage were grown in square pots (11 cm x 11 cm x 12 cm). All plants were kept in a glasshouse supplemented with artificial light (∼300 μmol m^-2^ s^-1^) with a 14h/10h light program, a temperature range of 22°C/18°C, and 40–60% relative humidity.

### Generation of maize mutants

Gene editing of *ZmCLE1E9* (Zm00001d003425), *ZmCLE1B3* (Zm00001d002849), *ZmBAM1A* (Zm00001d013162)/*ZmBAM1B* (Zm00001d034240) and *ZmBAM3C* (Zm00001d039218) were done with CRISPR-Cas9 technology. sgRNAs were designed by the CRISPR-P 2.0 web-tool based on the B73 reference genome (Liu *et al*, 2017). They were then synthesized and cloned into the pGW-Cas9 construct and transformed into the maize Hi-II hybrid background (A188 x B73) through *Agrobacterium*-mediated transformation (Char *et al*, 2017). Mutants were identified through sequencing the PCR amplicons from the targeted region. Cas9-negative plants were backcrossed to B73 for two generations to purify genetic background. Homozygous mutants and corresponding wildtype were selected from the BC2F2 population to propagate seeds for further experiments. Primers for gene editing and genotyping are listed in Table EV1.

### Plant treatment

Peptide treatment of detached plants or detached leaves were done by adding 5 µl 10 mM peptide stock solution in 12-ml tubes with 10 ml milli-Q water. Detached plants or leaves were taken from plants grown in pots, cut again under water with a razor blade, placed in a glass beaker filled with milli-Q water, incubated under dim light for at least 2 hours before use (Wang & Erb, 2024). Water treated plants were used as controls.

HAC treatment was done with volatile dispensers in glass cylinders supplied with clean airflow at a rate of 0.8 L min^-1^ as described before (Wang *et al*, 2023). Each HAC dispenser was prepared by adding 200 µl (Z)-3-hexenyl acetate in a 2 ml glass vial filled with 100 mg glass wool, sealed with a screw cap, pierced with a 1 µl glass capillary, sealed with PTFE tape and finally wrapped with aluminum foil. The emission rate was approximately 5 µg h^-1^, comparable to emission from herbivore infested maize plants (Mérey *et al*, 2011). No dispenser treatment was used as control.

Simulated herbivory was done by wounding the leaf blades 3 times with hemostatic forceps on each side of the mid-vein and adding 10 µl *Spodoptera exigua* oral secretions. Untreated plants were used as controls.

### Volatile measurements

Plant volatile profiling was conducted with a customized volatile sampling platform at room temperature as described before (Wang *et al*, 2023). All plants were measured inside of transparent glass cylinders (Ø 12 cm × H 45 cm) with a constant clear air flow(0.8 L min^-1^). Illumination during the experiments was provided by LED lights at an intensity of ∼300 μmol m^-2^ s^-1^ and a 14h/10h light program. For experiments with intact plants, the plants were transferred from the glasshouse to transparent glass cylinders for overnight acclimation. For experiments with detached seedlings, peptide treatment was done on the first day, 2 hours before dark; and HAC exposure was done on the 2^nd^ day, 2 hours after light was turned on. HAC dispensers stayed inside of the glass cylinders during the whole course of volatile profiling.

### Gene expression analysis

The 3rd leaves of V2 stage maize plants after HAC exposure were harvested and froze in liquid nitrogen for further analysis. Other than the HAC exposure time series experiments, plants were exposed to HAC for 2 hours. Total RNA extraction was done with the GeneJET PCR Purification Kit. DNAse I was used to remove genomic DNA from the RNA preparations before cDNA synthesis with a PrimeScript™ RT Master Mix. Gene expression levels were determined with the ORA™ SEE qPCR Green ROX L Mix kit and the QuantStudio 5 Real-Time PCR System. Relative gene expression was calculated after comparing to the expression of *ZmUBI1* (Zm00001d015327).

### Molecular docking of ZmBAM receptors and ZmCLE1E9 peptide

The structural model of the ZmBAM1A receptor was taken from the AlphaFold database (entry A0A1D6GGA2), models for ZmBAM1B and ZmBAM3C were obtained using AlphaFold server (https://alphafoldserver.com/). Briefly, the region comprehending the ectodomain (residues 21 to 621) was used for analysis and comparison with the Arabidopsis thaliana CLE-peptide receptor AtPXY in complex with its ligand AtCLE41 (PDB ID: 5GR9) (Zhang *et al*, 2016). Initial ZmCLE1E9 peptide orientation on the ZmBAM1A ectodomain was made based on the AtCLE41 position on the AtPXY/AtCLE41 complex. High resolution docking refinement and modelling was finally performed using the Roseta FlexPepDock Protocol (London *et al*, 2011). Images were generated using The PyMOL Molecular Graphics System, Version 2.0 Schrödinger, LLC.

### Stomatal phenotype determination

3D stomatal complex image acquisition was performed at the Stellaris5 Leica confocal microscope (Leica microsystems) with an excitation of 568 nm from a white light LED laser running at 85%. Images were acquired with a 63x glycerol objective (2-3x zoom, 2x line average, 1024 x 1024 pixels in stacks with a voxel size of 0.5 μm). Samples from the 2^nd^ leaves of V2 stage seedlings were cut at the 1/3 parts from the leaf tip, fixed in 12.5% acetic acid (2h), sequentially washed in 100% ethanol (4h), 50% ethanol (2h) and water (1h) before overnight staining in 0.02% Direct Red23 (in 1x PBS buffer). These samples were kept in PBS buffer until imaging on the same day (Zhou *et al*, 2023). Image analysis and processing were performed in Fiji (Schindelin *et al*, 2012).

Stomatal density, stomatal length and stomatal width were determined using nail polish imprint. The imprints were done by applying a thin layer of clear nail polish to the abaxial side of the 2^nd^ leaves (1/3 parts from the leaf tip) of V2 stage seedlings. These imprints were then observed at a 100-fold magnification using a Leica DM2500 optical231 microscope. Stomatal length was determined by the maximum length of guard cells. Stomatal width was determined by the maximum width of the subsidiary cells. Stomatal density was determined by counting the stomata in a 1.6 mm^2^ area.

Water uptake was calculated by measuring the remaining water compared to initial volume (10 mL) 24 hours after HAC exposure with a 14-h overnight peptide pretreatment. The cut stems were submerged in water during the whole experiment.

Stomatal conductance measurements were determined in a walk-in growth chamber (300 μmol m^-2^ s^-1^, 22°C, 60% RH) with a LI-COR LI-600 porometer. Stomatal conductance was determined 2 hours after peptide treatment on one side of the mid vein at the position of ∼15 cm from the tips of measured leaves. The 4^th^ and 5^th^ leaves of V4 stage B73 plants and 5^th^ and 6^th^ leaves of V5 stage *Zmbam1a1b3c* plants were used to guarantee full coverage of the 0.75 cm diameter aperture of the porometer.

### Statistical analysis

To understand the effects of treatment, genotype and/or time, we employed two sample t-tests or ANOVA. Upon significant ANOVA results, differences between specific groups were determined by multiple comparisons. Datasets that did not fit assumptions of ANOVA underwent log-transformation to meet requirements for equal variance and normality. For data that did not meet the assumptions of ANOVA even following transformation, differences were determined using non-parametric Kruskal-Wallis or Scheirer-Ray-Hare tests, in lieu of one- and two-way ANOVA, respectively. When results from non-parametric tests were significant, differences between specific groups were determined using pairwise Wilcoxan tests. See figure legends for details on statistical analyses.

## Data availability

All data used for generating figures in this study are available within the source data. Further information will be provided upon request to the corresponding authors.

## Author contributions

**Lei Wang**: Conceptualization; Funding acquisition; Investigation; Visualization; Methodology; Writing—original draft; Writing—review and editing. **Sara Hoefer**: Investigation. **Pedro Jimenez-Sandoval**: Investigation; Visualization. **Hao Yu**: Investigation. **Roxane Spiegelhalder:** Investigation. **Jamie Waterman**: Formal analysis. **Luke Hurni**: Investigation. **Lingfei Hu**: Investigation. **Lei Liu**: Resources. **David Jackson**: Resources. **Michael Raissig**: Resources. **Matthias Erb**: Conceptualization; Supervision; Funding acquisition; Writing—review and editing.

## Acknowledgements

We thank Sarah Dolder for excellent plant care. We also thank the whole Biotic Interactions group and Chemical Ecology group for helpful discussions. L.W. is funded by the Horizon 2020 Marie Skłodowska-Curie Actions (Grant Nr. 886651), the University of Bern and the Yazhouwan National Laboratory Fundamental Research Project (Grant Nr. 2310JM01). J.W. was funded by the Swiss National Science Foundation (Grant Nr. 210651). M.E. is funded by the European Research Council (ERC) under the European Union’s Horizon 2020 Research and Innovation Programme (ERC-2016-STG 714239), the Swiss National Science Foundation (Grant Nr. 200355), the Swiss State Secretariat for Education, Research, and Innovation (Project CANWAS) and the University of Bern.

